# Evaluation of MRI sequences for quantitative T1 brain mapping

**DOI:** 10.1101/195859

**Authors:** P Tsialios, M Thrippleton, C Pernet

## Abstract

T1 mapping constitutes a quantitative MRI technique finding significant application in brain imaging. It allows improved evaluation of contrast uptake, blood perfusion, volume, and provides more specific biomarkers of disease progression compared to conventional T1-weighted images. While there are many techniques for T1-mapping, there is also a wide range of reported T1-values in tissues, raising the issue of protocols’ reproducibility and standardization. The gold standard for obtaining T1-maps is based on acquiring IR-SE sequence. Widely used alternative sequences are IR-SE-EPI, VFA (DESPOT), DESPOT-HIFI and MP2RAGE that speed up scanning and fitting procedures. A custom MRI phantom was used to assess the reproducibility and accuracy of the different methods. All scans were performed using a 3T Siemens Prisma scanner. The acquired data were processed using two different codes. The main difference was observed for VFA (DESPOT) which grossly overestimated T1 relaxation time by 214 ms [CI: 126 270 ms] compared to the IR-SE sequence. MP2RAGE and DESPOT-HIFI sequences gave slightly shorter T1 than IR-SE (∼20 to 30ms) and can be considered as alternative and time-efficient methods for acquiring accurate T1 maps of the human brain, while IR-SE-EPI gave identical results, at a cost of a lower image quality.

## 1. Introduction

Among clinical examination modalities, Magnetic Resonance Imaging (MRI) is one of the most widely used, allowing to distinguish pathologic tissues with great precision. Contrast images like T1-weighted images are for instance used routinely as they preserve well anatomical details. These types of images, however, are qualitative images in the sense that they do not accurately measure tissue parameters such as T1 recovery time. Research efforts have shown interest in quantifying the longitudinal relaxation time (T1), especially in brain tissue [1] because of an increased sensitivity in pathology detection such as brain cancer, multiple sclerosis (integrity of myelin in the brain), ischemia, hepatic encephalopathy, or chronic alcoholism [2-5]. Moreover, quantitative T1 maps can help clinicians to evaluate contrast agent uptake, iron overload, and blood perfusion and volume [6]. They also provide a more robust template for morphometry studies and a more specific marker of disease progression in comparison to conventional T1 weighted images [7].

The gold standard method for T1 mapping makes use of inversion recovery spin echo (IR-SE) sequence [8,9]. However, IR-SE requires an extremely long scan time, which is impractical for clinical use and can lead to inaccurate measurements due to patient motion. Many efforts have been made to shorten the T1 measurement time, leading to a variety of techniques and a wide range of reported T1 values (from ∼1000 to 1600ms) raising the issue of protocols’ reproducibility and standardization [10]. The purpose of this study is to estimate the reproducibility and accuracy of T1 values acquired by four alternative T1 mapping protocols in a phantom and compare them to T1 values acquired by a gold standard protocol [6].

## 2. Methods

### 2.1 Imaging protocols

All imaging was performed at the University of Edinburgh on a 3 Tesla Siemens Prisma scanner using a custom MRI phantom [11]. The phantom (figure 1a) is filled with 1.5 g/l CuSO_4_ and 3.6 g/l NaCl solution and made of 9 tubes containing MnCl2 solution of various concentrations. Five different pulse sequences were used to image the phantom: a) IR-SE (TI = 30, 330 530, 780, 1030, 1530ms, TE/TR = 11/1550ms), b) IR-SE Echo Planar Imaging (IR-SE-EPI) (TI = 100, 340, 580, 820, 1060, 1300, 2000, 3000ms, TE/TR = 42/20000ms), c) Inversion Recovery Spoiled Gradient echo (IR-SPGR) or spoiled gradient echo (SPGR; TI = 600, 1500ms, TE/TR = 1,91/1170ms, flip angles = 2°, 5°, 12°), d) Variable Flip Angle (VFA; SPGR scans acquired in c) used) and e) Three Dimensional Magnetization Prepared Rapid Acquisition 2 Gradient Echoes (MP2RAGE) (TE/TR = 2.98/5000 ms). A thorough recording of phantom and room temperature was conducted in order to ensure the same experimental conditions as T1 increases with temperature by 2–3% per degree Celsius [12].

**Figure 1.**
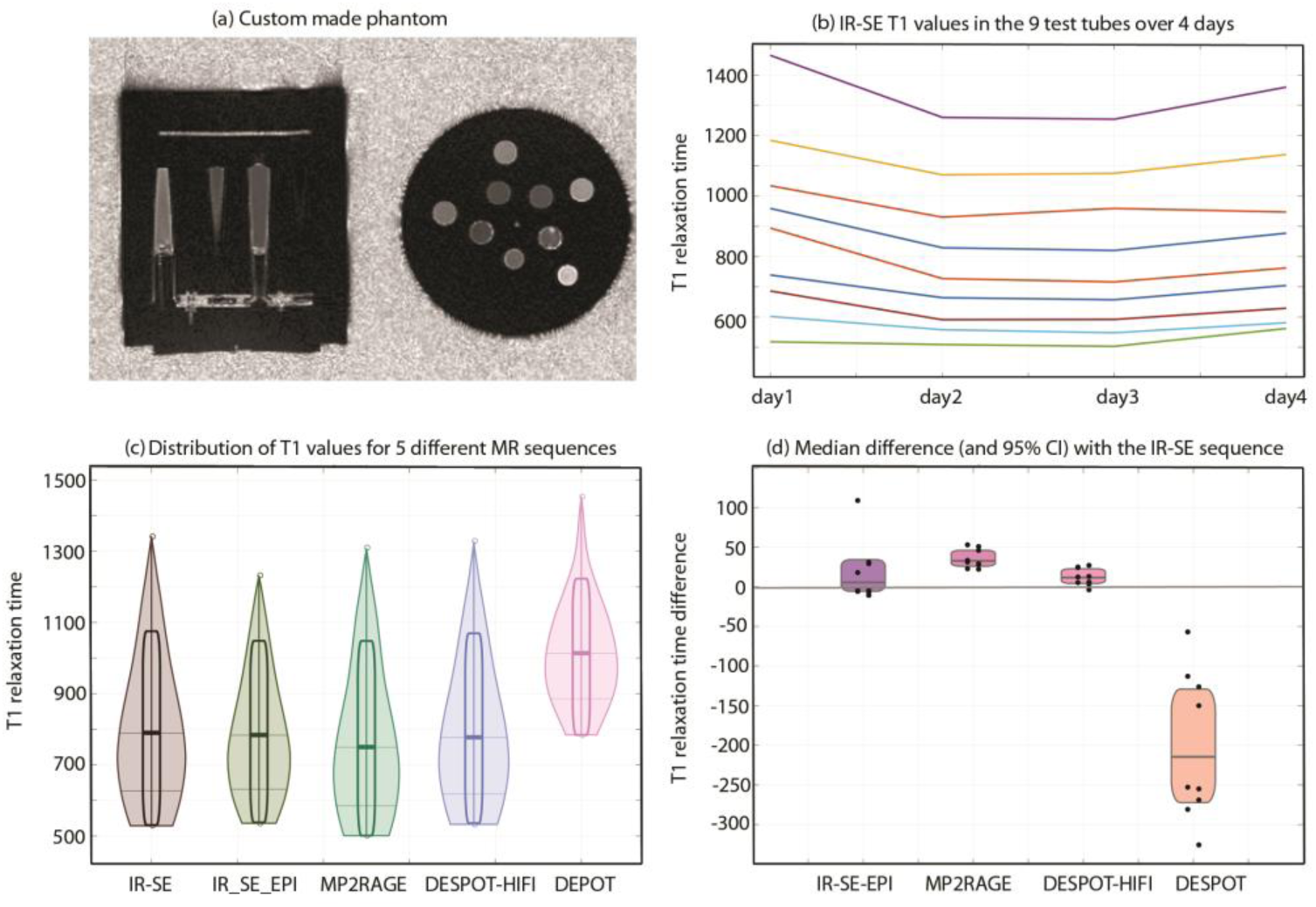
Panel (a) shows the MP2RAGE image of the phantom, panel (b) shows the T1 values acquired with IR-SE over 4 consecutive days, panel (c) shows the distributions of T1 values for the 9 test tubes using IR-SE, IR-SE-EPI, MP2RAGE, DESPOT-HIFI and VFA (DESPOT) acquired on the same day and panel (d), depicts the relative median differences from IR-SE-EPI, MP2RAGE, DESPOT-HIFI and VFA (DESPOT) to the gold standard IR-SE sequence. Violin plots show the non-parametric kernel of each data (Random Average Shifted Histograms) from the average ROI values and rectangles indicate the 95% confidence intervals of the medians (thick lines).

### 2.2 Image Processing

Image processing was carried out using two Matlab codes, the Stanford code (part of theT1Mapping Matlab package which was developed by Nikola Stikov and his colleagues at Stanford University [6]) and an in-house code (MJT) that uses a different method to fit the data. Both codes accept DICOM images where magnitude, phase, real and imaginary parts have been saved. Reported T1 values were obtained from each test tube by taking the average values across voxels for each TR (except MP2RAGE that provides directly the T1 value). Statistical comparisons were performed using a percentile bootstrap on median differences, with adjustment for multiple comparisons [13].

## 3 Results

The reproducibility of the gold standard IR-SE was evaluated by acquiring the sequence for four consecutive days (T1 values day1: 866 ms [531 1143], day2: 748 ms [509 1016] day3: 745 ms [511 1023] day4: 786 ms[564 1070], figure 1b). Small but significant fluctuations of T1 values were observed (p=.001) that might have risen from distributions in phantom temperature. Code comparison was performed on IR-SE and IR-SE-EPI sequences. Fitting procedures did not show difference on T1 values (2 ms [-6 13] and 4 ms [-9 9] respectively). Given the above results, comparison of sequences was performed from data acquired the same day and using MJT code (table 1). Significant differences were observed with VFA (DESPOT) showing much longer T1 values than other sequences (p=.001, +228 ms on average). DESPOT-HIFI (-11 ms [−22 −4] p=.002) and MP2RAGE (-32 ms [−45 −25] p=.001) had shorter values than the gold standard, with a significant difference between them (p=.001) (figure 1c and 1d).

**Table 1.** T1 values in ms (with median and 95% highest density interval) in each ROI.

| ROI | IR-SE | IR-SE-EPI | M2RAGE | DESPOT1-HIFI | VFA<br>(DESPOT1) |
| --- | --- | --- | --- | --- | --- |
| 1 | 872 | 843 | 850 | 866 | 1022 |
| 2 | 987 | 969 | 964 | 962 | 1044 |
| 3 | 1131 | 1100 | 1102 | 1104 | 1257 |
| 4 | 1341 | 1232 | 1310 | 1329 | 1454 |
| 5 | 529 | 536 | 501 | 533 | 784 |
| 6 | 582 | 588 | 549 | 577 | 835 |
| 7 | 631 | 642 | 585 | 618 | 900 |
| 8 | 702 | 706 | 649 | 678 | 983 |
| 9 | 763 | 767 | 712 | 760 | 1089 |
| median | 790 | 783 | 750 | 777 | 1013 |
| HDI | [530 1075] | [538 1048] | [501 1048] | [535 1069] | [784 1222] |

## 4 Discussion and Conclusion

The advantages of using quantitative imaging and especially T1 mapping methods in MRI studies are well-established. However, the main difficulty in gold-standard T1 mapping protocols is the impractically long scan time, creating the need for faster methods. Our results show that the gold standard IR-SE is relatively stable but some values seems more sensitive to temperature variations than others, b) data fitting procedure (when properly implemented) have little impact on T1 values c) VFA (DESPOT1) T1 maps give longer T1 values on our scanner in comparison to the other methods, d) MP2RAGE and DESPOT-HIFI T1 values are close to the gold standard but still differ significantly.

## Acknowledgement

The authors would like to thank Nikola Stikov, Assistant Professor of Biomedical Engineering at Polytechnique Montréal for providing them the “T1Mapping Matlab package”. Moreover, part of this work was partially supported by European Union resources under the framework of the TRIMAGE: “A dedicated trimodality (PET/MR/EEG) imaging tool for schizophrenia” project.

